# Experimental and *In Silico* interaction studies of Alpha Amylase-Silver nanoparticle: a nano-bio-conjugate

**DOI:** 10.1101/2022.06.11.495728

**Authors:** Awadhesh Kumar Verma, Abhijeet Mishra, Tarun Kumar Dhiman, Meryam Sardar, Pratima R. Solanki

**Affiliations:** Special Centre for Nanoscience, Jawaharlal Nehru University, New Delhi,110067, India; Department of Biosciences, Jamia Millia Islamia University, New Delhi,110025, India

**Keywords:** Ag NPs, Alpha amylase, *In silico*, Docking, Ag-Nanocluster

## Abstract

In the current work, biosynthesis of silver nanoparticle (Ag NPs) and interaction study between alpha amylase and Ag NPs/nanocluster has been performed *via* wet-lab as well as *in silico* approach. We have synthesized Ag NPs using alpha amylase enzyme which reduces the silver nitrate precursor forming the stable Ag NPs. UV-Visible spectroscopy and fluorescence spectroscopies were performed for optical characterization of Ag NPs. UV-Vis spectra showed the wide absorption band centered around 475 nm due to surface plasmon resonance. We have also observed gradual decrease in fluorescence intensity with the increase in incubation time. Also, shift in λmax of the emission spectra was recorded which clearly suggested the formation of nano-bio-conjugate. Circular dichroism spectra show the initial decrease in the ellipticity, when we added the silver nitrate, but after incubating for different time, there are no major changes in secondary structure of protein. In computational study we have modelled ground state configuration of (Ag)_24_ nanocluster using *in silico* approach. Further docking of the modelled optimized nanocluster with alpha amylase was performed and found that Ag-nanocluster showing non-covalent interaction with alpha amylase and forming stable docking complex.

## 1. Introduction

Understanding the bio-molecular interactions between any enzyme and its respective substrate or any different molecules like its inhibitors, activators, or any drug molecules are crucial for life science research.[1–5] In recent days especially the nano-bio-molecular interaction has attracted the researcher and scientist in the field of nano-biotechnology all across the globe. As the nanobiotechnology field has promising future prospect for the improvement of complete healthcare management for the patient begin with diagnosis up to the treatment.[6–10] It will make us enable for rapid testing as well as the early-stage diagnosis of the diseases [11–13]. Silver nanoparticles (Ag NPs) are most widely used in textile industries, cosmetics, water treatment, antimicrobial drugs, antioxidant and antidiabetic activity etc. [14–16] It is mostly preferred due to its specialized magnetic, electrical and optical properties. These properties of nanomaterials not only dependent on the shape and size of the nanoparticle but also the different route of synthesis[17,18] As far as the chemical methods are concerned it applies the reduction process of precursor metal salts by the chemical reagents like polyvinyl pyrrolidine, hydroxylamine, sodium borohydride etc.[19,20] It is very much true that nanomaterials produced by these methods are having maximum purity as well as well-defined properties, However, the nanomaterial synthesized through this process may not be suitable for biochemical and biological applications due to toxicity and biocompatibility issues.[21–24] To overcome these challenges, the biosynthetic process may be one of the alternate ways which cares not only about toxicity but also the biocompatibility and is environmentally suitable. [25–27]

In recent years’ huge number of microbes have been involved in the production of metal nanomaterials however the mechanism of biosynthetic route for nanomaterial synthesis has not been established yet.[28,29] Phytochelatin and NADPH dependent nitrate reductase based *in vitro* synthesis of Ag NPs stabilized through peptide capping and having size 10-25 nm in diameter have also been reported in the research article. [30] Extracellular biosynthesis of silver nanomaterials using *Aspergillus flavus* NJP08 was reported by Jain et al., and they confirmed the presence of extracellular protein through UV-Visible and FTIR spectroscopy. [31] The presence of 2 bands of protein one 32 kDa and 35 kDa in SDS-PAGE proved that these are mainly responsible for the synthesis and stability of Ag NPs. They also predicted that protein component 32 kDa might be a reductase secreted by the fungus.

Computational docking which often called as *in silico* molecular docking or merely docking is a computational approach, that strives for mimicking the binding of ligand molecule to the protein molecule.[32–36] It predicts the optimum binding conformation. Orientation of molecules interacting with each other in 3D space including estimation of binding energy, binding affinity or a scoring function that represents the strength of binding, 3D structure and stability of the protein-ligand complex.[37–40] It gives the detailed information about the binding of ligand molecule in active sites, orientation of ligand molecule, which conformation of ligand is most stable in the binding site and leads to the *de novo* design and optimization of the ligand. [41,42] It provides multiple analyses based on diverse scoring methods and selection criteria for bindings[43–45] It utilizes the idea of complementarity of the molecules in which the structures interact with each other such as hands in a glove, in which both physico-chemical properties and shape of structures bestow best fitting among molecules.[46,47] It predicts the orientation and positions of the molecule in the binding site. In modern drug discovery, it is considered as an indispensable constituent.[48,49] In modern era, most of the biotech and pharmaceutical companies have been using this approach most extensively and routinely for wide range of applications. [50] In computational docking most of the docking software uses search algorithm which generates a large number of poses of molecules in the binding sites and scoring functions which calculate perfect score or binding affinity accurately for a particular pose.[51] Although the computational docking has some technical limitations however, overall docking is good enough. Here, we have used AutoDock Vina as docking software. [52] In current work, the biosynthesis and interaction study between alpha amylase and Ag NPs/nano cluster has been performed *via* both wet lab as well as *in silico* approach.

## 2. Materials and Methods

Pure Alpha-amylase was purchased from Hi-media Lab Pvt. Ltd., India and Tris-buffer, silver nitrate, were bought from Merck Ltd., India Tris-HCl buffer was prepared by mixing of 20 mM Tris with 6 mM NaCl by maintaining pH 8.0. 1 mM aqueous solution of silver nitrate was prepared using MilliQ water in the dark bottle as it is sensitive for light. 2 mg/ml alpha amylase enzyme solution was made in Tris-HCl buffer at pH 8.0. All the materials were used in the proper ratio and reaction was performed in the proper way.

### 2.1 Biosynthesis and purification of Ag NPs

Biosynthesis of the Ag NPs was performed by incubation of 40 ml alpha-amylase solution (having 2 mg/ml concentration in Tris-HCl buffer at pH 8.0) and 60 ml of 1 mM freshly prepared silver nitrate solution in MilliQ water. [53,54] The reaction mixture was kept at 30°C and pale brown color of solution after few minutes was seen, distinctly visible from fresh AgNO_3_ solution which shows the synthesis of Ag NPs.[55] After successive time intervals the aliquots were taken out and the synthesis of Ag NPs was monitored by UV-VIS spectroscopy. Synthesized Ag NPs solution was subjected to centrifugation process at 9,000 rpm at 4°C for 5 minute and supernatant was discarded. The entire pellet was washed four times in distilled water than air dried; which can be further used for the characterization studies.

### 2.2 Computational Details

For the structure modeling, first we generated the structure of (Ag)_24_ nanocluster by drawing it on Marvin sketch followed by 2D and 3D cleaning every time and checking its structure by visualizing in Marvin view for 3D geometrical conformation. (Figures are shown in results section). Further, we calculated each and every parameter like atomic co-ordinates (XYZ), bond length and bond angle using ChemBioDraw Ultra software and optimized the structure. Alpha amylase structure was obtained from Protein Dada Bank (PDB) having (PDB ID: 3VX1) crystal structure of alpha-amylase from *Aspergillus oryzae* obtained from RCSB-PDB.[56] We performed rigid docking of Ag-nanocluster with alpha amylase using AuotoDockVina to get the insight of nano-bio interaction. [57] PyMol, Biovia Discovery Studio and VMD software have been utilized for the identification and visualization of the residue in the active site. [58–60]

## 3. Results and Discussion

### 3.1 Optical Studies

The UV-Visible spectra of the solution mixture were recorded at regular time intervals consisting of an increasing intense absorption band at 475 nm, the absorption increased up to 36 hours and after that it was stable (as shown in Figure 2. (a)). [61, 62] An absorption band at 260– 280 nm was clearly visible due to the enzyme alpha amylase. The protein band also increased with time. This increase in absorbance with time had been reported earlier also. The control sample not showing any kind of color and did not have any kind of peak in the visible range confirming that alpha amylase is mainly responsible for Ag NPs synthesis.

**Fig. 1:**
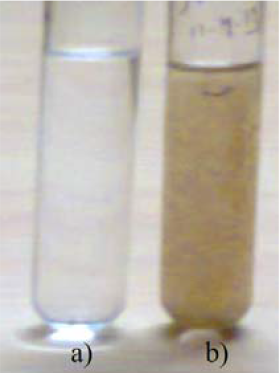
Visual observation of the solution (a) before and (b) after Ag NPs formation. The enzyme solution having concentration 2 mg/ml in Tris-HCl buffer at pH 8.0 and 1 mM AgNO_3_ in MilliQ water at the beginning of the reaction (1st tube) and after few hours of mixing (2nd tube).

**Fig. 2:**
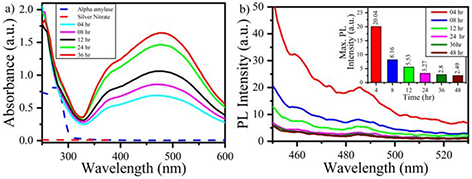
(a) shows UV-Visible absorption spectra recorded at regular interval of time the biosynthesis of Ag NPs mediated by alpha-amylase at 30°C, (b) shows fluorescence spectra recorded at regular interval of time the biosynthesis of Ag NPs mediated by alpha-amylase at 30°C.

Fluorescence spectra were recorded on a Cary Eclipse Fluorescence Spectrophotometer using 1cm quartz cell. A slit width of 5 nm was used for both excitation and emission. Samples were excited at 420 nm and the emission spectra were recorded ranging from 440 nm to 540 nm using a scanning speed 600 nm / minute at 750 volts at 25°C. From the above spectra as shown in Figure 2. (b), it is clear that the fluorescence intensity of mixture solution is gradually decreased with the increase of time which shows that there is quenching of fluorescence intensity with the increasing time of reaction-mixture so nano-bio interaction occurred between alpha amylase and Ag NPs.

To assess the involvement of tryptophan during synthesis of Ag NPs; fluorescence spectra of tryptophan was recorded. For this same set of experiment was performed under similar conditions and same samples were excited at 295 nm and the respective emission spectra were recorded ranging from 310 to 400 nm using a scanning speed 600 nm/min at 750 volts at 25°C. From the above spectra [Figure 3 (a) and (b)], it is clear that there is gradual decrement in fluorescence intensity of tryptophan present in alpha amylase, with the increase in time of incubation after the addition of sliver nitrate and also there was a shift in λmax of the emission spectra as shown in Figure 4. (a).

**Fig. 3:**
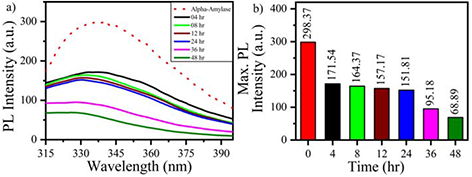
(a) Fluorescence spectra of alpha amylase in the absence and presence of silver nitrate solution at different time interval for enzymatic synthesis as well as nano-bio conjugation of silver nanoparticles-alpha-amylase.

**Fig.4:**
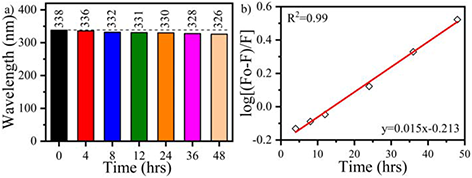
a shows graphs of alpha amylase Trp fluorescence in Ag-alpha amylase system as a function of time. The system shows the decrease of Trp fluorescence as a function of incubation time. The excitation energy was 295 nm and emission spectra obtained from 290-400 nm.

**Fig.5:**
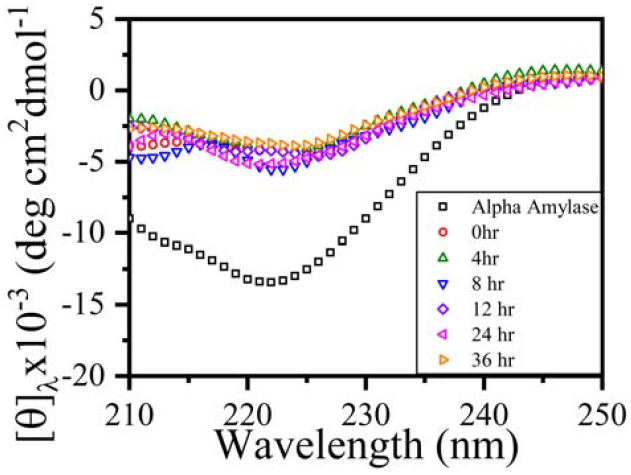
CD spectra of reaction mixture recorded at regular interval of time for the biosynthesis of Ag NPs as well as to monitor the nano-bio-interaction. Initially there was decrement in the ellipticity but after incubating silver nitrate precursor solution for different time interval we observed that, there is no change in ellipticity. It means that incubation of AgNO_3_ not affecting the structure so might be possible that there is no major change in secondary structure of alpha amylase during synthesis of silver nanoparticles and nano-bio-conjugation.

Figure 4. (b) shows the log [(Fo-F)/F] vs. [time] graph showing linearity in quenching of fluorophores accessible to the quencher. Figure 3. (a) and (b) and Figure 4. (a) and (b) clearly suggesting the formation of nano-bio-conjugate. It might be due to the exposure of tryptophan residue on the outer surface of protein, which may have participated during the nano-bioconjugate formation.

### 3.2 Circular Dichroism study

Circular Dichroism (CD) studies were carried out to observe the effect of Ag NPs conjugation on secondary and tertiary structure of Alpha-Amylase. CD spectra were obtained using an AVIV 420SF CD Spectrophotometer. The instrument routinely went under calibration using D-10 camphor sulfonic acid and purging of N_2_ was done continuously to the lamp, optics and chamber of sample in 1:3:1 ratio. The main function of purging the CD spectrophotometer with nitrogen gas is to removing oxygen from the housing lamp, monochromator, as well as the chamber of sample. If there is a presence of oxygen, it may to the ozone formation causing optics degradation. Another reason for removal of oxygen is that it absorbs deep UV photons, hence will reduce availability of light for measurement purpose. Far-UV CD spectra (206-250 nm) were recorded with a cell of 1 mm path length at temperature 25°C, bandwidth 1nm, wavelength start at 250 nm and end at 206 nm, having wavelength step 1nm and averaging time 5 second. For making blank contribution each spectrum was corrected. Each spectrum was average of 3 scans with the average time of 1 sec/nm. CD data was analyzed by online available software, K2d.[63]

### 3.3 *In silico* Approach for Modelling and Docking Studies on Alpha amylase-(Ag)_24_ Complex Theoretical Modeling of (Ag)_24_ Nanocluster

3D cage like (Ag)_24_ nanocluster in ground state configuration was theoretically modeled (Figure. 6) using Marvin Sketch and ChemBioDraw ultra software which utilizes DFT (density function theory) approach. Figure.6 (a) shows the ground state configuration (Ag)_24_ nano cluster as space filling model. The grey color ball represents the Ag atoms present in the nanocluster. Ground state configuration (Ag)_24_ nanocluster as space filling model having 8 hexagonal and 4 square rings. Figure 6. (b) shows the 3D surface view of (Ag)_24_ nanocluster (Connolly molecular) along with the Cartesian coordinates shown in tabular form, right side of the Figure 6, generated from the structure drawn after optimization and energy minimization.

**Figure 6.**
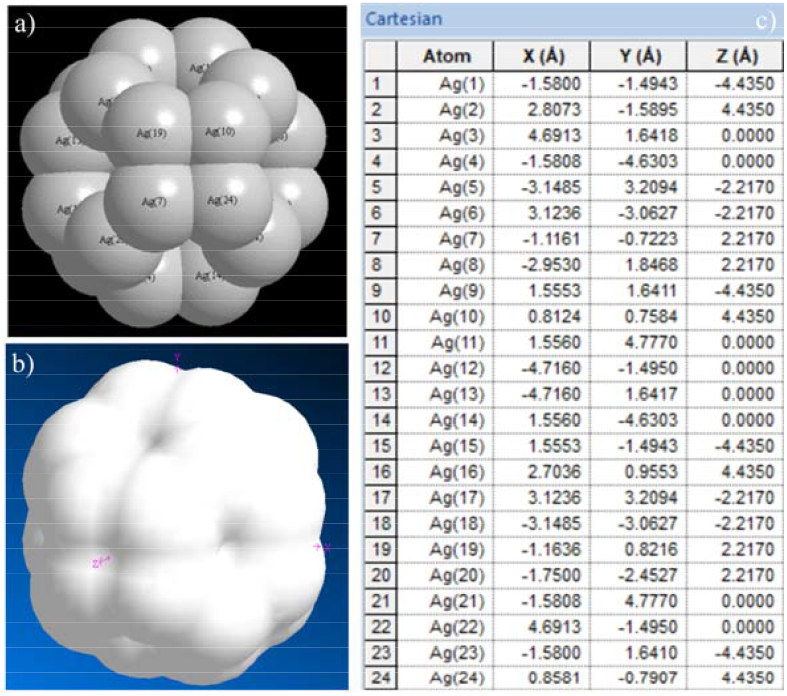
(a) and (b) 3D ground state configuration (Ag)_24_ nanocluster (space filling model along with coordinates view) made by using Marvin Sketch and ChemBioDraw Ultra software which utilizes DFT (Density Function Theory) approach, and (c) Ground state configurations (Ag)_2_4 nanocluster showing surface view (Connolly molecular) in 3D space along with X, Y & Z coordinate axes.

The distance between two Ag atoms is 2.6682 to 2.6894 Å, as shown in Table 1. The bond angle ranging between 89° to 120°.

**Table 1:** shows the bond length (in Å) bond angle and dihedral angle of all possible confirmation of atoms of Ag-nanocluster in 3D space.

| Sr. No. | Atom | Bond Atom | Bond length (Å) | Angle Atom | Bond Angle (°) | 2<sup>nd</sup> Angle Atom | 2<sup>nd</sup> Angle (°) | Conformation |
| --- | --- | --- | --- | --- | --- | --- | --- | --- |
| 1 | Ag (1) |  |  |  |  |  |  |  |
| 2 | Ag (15) | Ag (1) | 2.685 |  |  |  |  |  |
| 3 | Ag (6) | Ag (15) | 2.6679 | Ag (1) | 120.0003 |  |  |  |
| 4 | Ag (9) | Ag (15) | 2.685 | Ag (1) | 90.0048 | Ag (6) | 120.0037 | Pro-R |
| 5 | Ag (14) | Ag (6) | 2.685 | Ag (15) | 120 | Ag (1) | -0.0018 | Dihedral |
| 6 | Ag (22) | Ag (6) | 2.685 | Ag (14) | 90.0008 | Ag (15) | 119.9989 | Pro-S |
| 7 | Ag (2) | Ag (22) | 2.685 | Ag (6) | 89.9991 | Ag (14) | 0.0001 | Dihedral |
| 8 | Ag (3) | Ag (22) | 2.6678 | Ag (2) | 120.0007 | Ag (6) | 119.9997 | Pro-S |
| 9 | Ag (4) | Ag (14) | 2.6678 | Ag (2) | 119.9999 | Ag (6) | 120.0001 | Pro-R |
| 10 | Ag (24) | Ag (2) | 2.6678 | Ag (14) | 120.0006 | Ag (22) | 119.999 | Pro-S |
| 11 | Ag (7) | Ag (24) | 2.685 | Ag (2) | 119.9998 | Ag (14) | -0.0014 | Dihedral |
| 12 | Ag (10) | Ag (24) | 2.685 | Ag (2) | 120.001 | Ag (7) | 89.9975 | Pro-R |
| 13 | Ag (16) | Ag (3) | 2.685 | Ag (22) | 119.9995 | Ag (6) | 109.4705 | Dihedral |
| 14 | Ag (11) | Ag (16) | 2.685 | Ag (3) | 89.9992 | Ag (10) | 120.0015 | Pro-R |
| 15 | Ag (17) | Ag (9) | 2.6679 | Ag (15) | 119.9957 | Ag (1) | 125.2665 | Dihedral |
| 16 | Ag (19) | Ag (10) | 2.685 | Ag (16) | 119.9986 | Ag (24) | 90.0026 | Pro-R |
| 17 | Ag (8) | Ag (19) | 2.6679 | Ag (7) | 120.0015 | Ag (10) | 120.0007 | Pro-S |
| 18 | Ag (20) | Ag (4) | 2.685 | Ag (14) | 119.9993 | Ag (2) | 0.0011 | Dihedral |
| 19 | Ag (12) | Ag (20) | 2.685 | Ag (4) | 90.0007 | Ag (7) | 120.0002 | Pro-S |
| 20 | Ag (18) | Ag (4) | 2.685 | Ag (14) | 119.9994 | Ag (20) | 89.9995 | Pro-S |
| 21 | Ag (13) | Ag (12) | 2.6679 | Ag (18) | 119.9995 | Ag (20) | 119.9993 | Pro-S |
| 22 | Ag (5) | Ag (13) | 2.685 | Ag (8) | 90.0006 | Ag (12) | 120 | Pro-R |
| 23 | Ag (21) | Ag (11) | 2.6678 | Ag (16) | 119.9979 | Ag (17) | 120.0099 | Pro-S |
| 24 | Ag (23) | Ag (9) | 2.685 | Ag (15) | 89.9986 | Ag (17) | 119.9994 | Pro-R |

### 3.4 Docking of (Ag)_24_ nanocluster with alpha amylase

In the current work we have also utilized computational approaches like molecular docking which could provide the molecular insights into the interaction between Ag nanocluster and alpha amylase which may not be accessible from experiments alone. For this, we did theoretical modelling of 3D cage like silver nanocluster in ground state configuration using Marvin sketch software. We further performed rigid docking of (Ag)_24_ nanocluster with alpha amylase (PDB ID: 3VX1. crystal structure of alpha-amylase from *Aspergillus oryzae* obtained from RCSB-PDB) using AuotoDockVina to get the insight of nano-bio interaction. Alpha-amylase has total 478 amino acid residues, among these only 12 residues are tryptophan. Figure.7 shows the docked structure of Ag-nanocluster with alpha amylase drawn using pymol software in which some part of the amino acid residue is showing involvement of non-covalent interaction with Ag-nanocluster and it clearly describes that Ag-nanocluster is bound to the protein surface in its active site. The detail information for bond length, bond angle and dihedral angle for each atom of Ag nanocluster has been shared in supplementary data.

**Fig.7:**
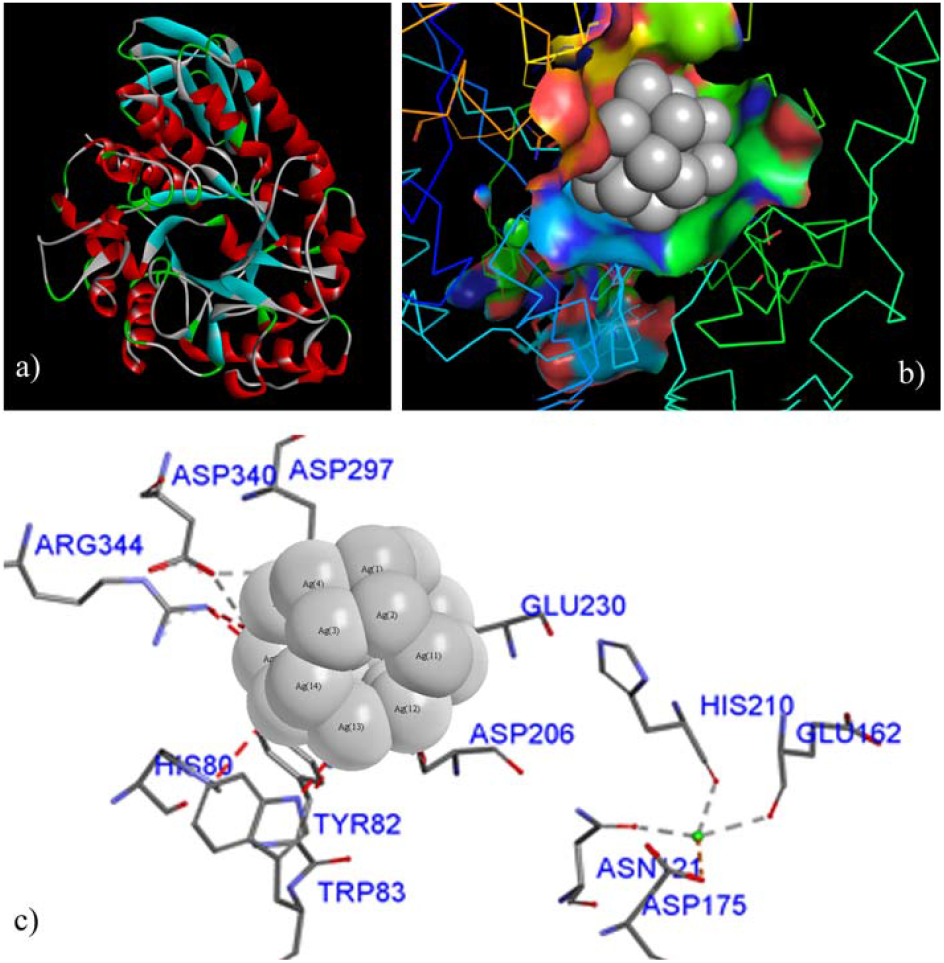
(a) shows the docked structure of (Ag)_24_-nanocluster with alpha amylase, (b) shows involvement of non-covalent interaction with Ag-nanocluster drawn using pymol software, and (c) shows the interactions predicted by the docking study. HIS 80, TRP 83, TYR 82, ASN 121, GLU 162, ASP 206, HIS 210, GLU 230, ASP 297, ASP 340, ARG 344 were observed to be involved in interaction with (Ag)_24_ nano cluster. From docking result shown in figure, it is clear that the (Ag)24 nano cluster is binding with alpha amylase.

## 4. Conclusion

We adopted the above method of nanomaterial synthesis as it is considered as ecologically safe and sound, an alternative way over conventional route of physical and chemical mode of synthesis. The gradual decrease in fluorescence intensity with the increase in time of incubation and also there was a shift in λmax of the emission spectra which clearly suggested the formation of nano-bio-conjugate. It might be due to the exposure of tryptophan residue on the outer surface of protein, which may have participated during the nano-bio-conjugate formation. From our computational study, we observed that Ag NPs/cluster interacted with alpha amylase non-covalently and forming a stable nano-bio conjugate. So, after performing all the necessary analysis it can be concluded that the developed nano-bio conjugate may be used in the development of nanosensor based on Ag-Alpha amylase nano bioconjugate which can be used potentially in food and environmental safety applications.

## Acknowledgments

The author would highly acknowledge Prof. B. Jayram and Mr. Shashank Shekhar to provide the computational facility at SCF-Bio IIT Delhi and Prof. Shashank Deep Department of Chemistry, Indian Institute of Technology, Delhi to provide the experimental facility and guidance for research work.

---

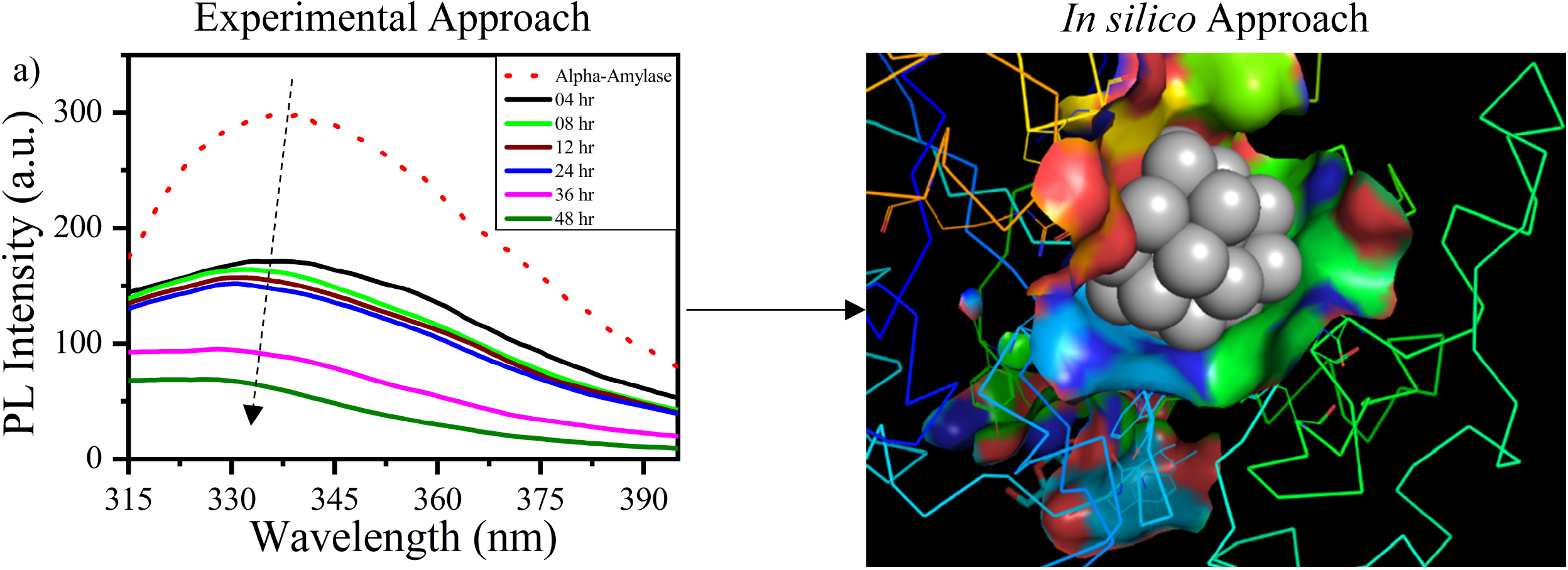

## Notes

### Competing Interest Statement

The authors have declared no competing interest.

