## Supplementary File for "Experimental and *In Silico* interaction studies of Alpha Amylase-Silver nanoparticle: a nano-bio-conjugate"

**Table 1:** Ground state configurations (Ag)_24_ nano cluster showing spacial arrangement of Ag atoms with bond length.

| **Atoms** | **Bond length (Å)** |
| --- | --- |
| Ag(2)-Ag(22) | 2.685 |
| Ag(16)-Ag(3) | 2.685 |
| Ag(2)-Ag(14) | 2.685 |
| Ag(7)-Ag(24) | 2.685 |
| Ag(7)-Ag(20) | 2.6678 |
| Ag(7)-Ag(19) | 2.685 |
| Ag(20)-Ag(4) | 2.685 |
| Ag(12)-Ag(20) | 2.685 |
| Ag(8)-Ag(19) | 2.6678 |
| Ag(8)-Ag(21) | 2.685 |
| Ag(13)-Ag(8) | 2.685 |
| Ag(11)-Ag(16) | 2.685 |
| Ag(18)-Ag(12) | 2.685 |
| Ag(12)-Ag(13) | 2.6678 |
| Ag(11)-Ag(21) | 2.6678 |
| Ag(17)-Ag(11) | 2.685 |
| Ag(10)-Ag(24) | 2.685 |
| Ag(19)-Ag(10) | 2.685 |
| Ag(16)-Ag(10) | 2.6678 |
| Ag(23)-Ag(9) | 2.685 |
| Ag(17)-Ag(9) | 2.6679 |
| Ag(15)-Ag(9) | 2.685 |
| Ag(6)-Ag(22) | 2.685 |
| Ag(6)-Ag(15) | 2.6678 |
| Ag(6)-Ag(14) | 2.685 |
| Ag(5)-Ag(23) | 2.6678 |
| Ag(21)-Ag(5) | 2.685 |
| Ag(13)-Ag(5) | 2.685 |
| Ag(18)-Ag(4) | 2.685 |
| Ag(4)-Ag(14) | 2.6678 |
| Ag(3)-Ag(22) | 2.6678 |
| Ag(3)-Ag(17) | 2.685 |
| Ag(24)-Ag(2) | 2.6679 |
| Ag(23)-Ag(1) | 2.685 |
| Ag(1)-Ag(18) | 2.6678 |
| Ag(15)-Ag(1) | 2.685 |

**Table 2:** Ground state configurations (Ag)_24_ nano cluster showing spacial arrangement of Ag atoms with bond angle.

| **Bond Angle** | **Degree (º)** |
| --- | --- |
| Ag(7)-Ag(24)-Ag(10) | 89.9953 |
| Ag(7)-Ag(24)-Ag(2) | 119.9929 |
| Ag(10)-Ag(24)-Ag(2) | 119.9955 |
| Ag(9)-Ag(23)-Ag(5) | 119.9948 |
| Ag(9)-Ag(23)-Ag(1) | 90.0045 |
| Ag(5)-Ag(23)-Ag(1) | 119.9942 |
| Ag(2)-Ag(22)-Ag(6) | 89.9979 |
| Ag(2)-Ag(22)-Ag(3) | 119.994 |
| Ag(6)-Ag(22)-Ag(3) | 119.9987 |
| Ag(8)-Ag(21)-Ag(11) | 120.0025 |
| Ag(8)-Ag(21)-Ag(5) | 90.0029 |
| Ag(11)-Ag(21)-Ag(5) | 119.9923 |
| Ag(7)-Ag(20)-Ag(4) | 119.9964 |
| Ag(7)-Ag(20)-Ag(12) | 120.0062 |
| Ag(4)-Ag(20)-Ag(12) | 90.0029 |
| Ag(7)-Ag(19)-Ag(8) | 120.008 |
| Ag(7)-Ag(19)-Ag(10) | 89.9956 |
| Ag(8)-Ag(19)-Ag(10) | 120.003 |
| Ag(12)-Ag(18)-Ag(4) | 90.0029 |
| Ag(12)-Ag(18)-Ag(1) | 119.9957 |
| Ag(4)-Ag(18)-Ag(1) | 119.9993 |
| Ag(11)-Ag(17)-Ag(9) | 119.9903 |
| Ag(11)-Ag(17)-Ag(3) | 89.9982 |
| Ag(9)-Ag(17)-Ag(3) | 120.0018 |
| Ag(3)-Ag(16)-Ag(11) | 89.9983 |
| Ag(3)-Ag(16)-Ag(10) | 119.9947 |
| Ag(11)-Ag(16)-Ag(10) | 120.0022 |
| Ag(9)-Ag(15)-Ag(6) | 120.0037 |
| Ag(9)-Ag(15)-Ag(1) | 90.0048 |
| Ag(6)-Ag(15)-Ag(1) | 119.997 |
| Ag(2)-Ag(14)-Ag(6) | 89.9977 |
| Ag(2)-Ag(14)-Ag(4) | 119.9929 |
| Ag(6)-Ag(14)-Ag(4) | 119.9966 |
| Ag(8)-Ag(13)-Ag(12) | 120.0092 |
| Ag(8)-Ag(13)-Ag(5) | 90.003 |
| Ag(12)-Ag(13)-Ag(5) | 119.9963 |
| Ag(20)-Ag(12)-Ag(18) | 89.9971 |
| Ag(20)-Ag(12)-Ag(13) | 119.9924 |
| Ag(18)-Ag(12)-Ag(13) | 120.0035 |
| Ag(16)-Ag(11)-Ag(21) | 119.9979 |
| Ag(16)-Ag(11)-Ag(17) | 90.0017 |
| Ag(21)-Ag(11)-Ag(17) | 120.0099 |
| Ag(24)-Ag(10)-Ag(19) | 90.0046 |
| Ag(24)-Ag(10)-Ag(16) | 120.0044 |
| Ag(19)-Ag(10)-Ag(16) | 119.9974 |
| Ag(23)-Ag(9)-Ag(17) | 120.0073 |
| Ag(23)-Ag(9)-Ag(15) | 89.9953 |
| Ag(17)-Ag(9)-Ag(15) | 119.9957 |
| Ag(19)-Ag(8)-Ag(21) | 119.997 |
| Ag(19)-Ag(8)-Ag(13) | 119.9906 |
| Ag(21)-Ag(8)-Ag(13) | 89.9969 |
| Ag(24)-Ag(7)-Ag(20) | 120.0046 |
| Ag(24)-Ag(7)-Ag(19) | 90.0045 |
| Ag(20)-Ag(7)-Ag(19) | 119.9936 |
| Ag(22)-Ag(6)-Ag(15) | 119.9991 |
| Ag(22)-Ag(6)-Ag(14) | 90.0022 |
| Ag(15)-Ag(6)-Ag(14) | 120.0041 |
| Ag(23)-Ag(5)-Ag(21) | 120.0054 |
| Ag(23)-Ag(5)-Ag(13) | 120.0049 |
| Ag(21)-Ag(5)-Ag(13) | 89.9971 |
| Ag(20)-Ag(4)-Ag(18) | 89.9971 |
| Ag(20)-Ag(4)-Ag(14) | 120.0051 |
| Ag(18)-Ag(4)-Ag(14) | 120.0018 |
| Ag(16)-Ag(3)-Ag(22) | 120.0063 |
| Ag(16)-Ag(3)-Ag(17) | 90.0018 |
| Ag(22)-Ag(3)-Ag(17) | 120.001 |
| Ag(22)-Ag(2)-Ag(14) | 90.0022 |
| Ag(22)-Ag(2)-Ag(24) | 120.0051 |
| Ag(14)-Ag(2)-Ag(24) | 120.0081 |
| Ag(23)-Ag(1)-Ag(18) | 120.0053 |
| Ag(23)-Ag(1)-Ag(15) | 89.9954 |
| Ag(18)-Ag(1)-Ag(15) | 120.0013 |

**Table 3:** Ground state configurations (Ag)_24_ nano cluster showing spacial arrangement of Ag atoms with dihedral angle.

| **Dihedral Angle** | **Degree (º)** |
| --- | --- |
| Ag(24)-Ag(2)-Ag(22)-Ag(3) | -0.0111 |
| Ag(24)-Ag(2)-Ag(22)-Ag(6) | -125.2699 |
| Ag(14)-Ag(2)-Ag(22)-Ag(3) | 125.2669 |
| Ag(14)-Ag(2)-Ag(22)-Ag(6) | 0.008 |
| Ag(10)-Ag(16)-Ag(3)-Ag(17) | 125.2703 |
| Ag(10)-Ag(16)-Ag(3)-Ag(22) | 0.0009 |
| Ag(11)-Ag(16)-Ag(3)-Ag(17) | 0.0066 |
| Ag(11)-Ag(16)-Ag(3)-Ag(22) | -125.2629 |
| Ag(24)-Ag(2)-Ag(14)-Ag(4) | 0.0118 |
| Ag(24)-Ag(2)-Ag(14)-Ag(6) | 125.2675 |
| Ag(22)-Ag(2)-Ag(14)-Ag(4) | -125.2637 |
| Ag(22)-Ag(2)-Ag(14)-Ag(6) | -0.008 |
| Ag(19)-Ag(7)-Ag(24)-Ag(2) | -125.2556 |
| Ag(19)-Ag(7)-Ag(24)-Ag(10) | -0.0029 |
| Ag(20)-Ag(7)-Ag(24)-Ag(2) | 0.006 |
| Ag(20)-Ag(7)-Ag(24)-Ag(10) | 125.2587 |
| Ag(19)-Ag(7)-Ag(20)-Ag(12) | 0.0024 |
| Ag(19)-Ag(7)-Ag(20)-Ag(4) | 109.4799 |
| Ag(24)-Ag(7)-Ag(20)-Ag(12) | -109.4738 |
| Ag(24)-Ag(7)-Ag(20)-Ag(4) | 0.0037 |
| Ag(20)-Ag(7)-Ag(19)-Ag(10) | -125.2678 |
| Ag(20)-Ag(7)-Ag(19)-Ag(8) | 0.0004 |
| Ag(24)-Ag(7)-Ag(19)-Ag(10) | 0.0029 |
| Ag(24)-Ag(7)-Ag(19)-Ag(8) | 125.271 |
| Ag(12)-Ag(20)-Ag(4)-Ag(14) | 125.2669 |
| Ag(12)-Ag(20)-Ag(4)-Ag(18) | 0.0004 |
| Ag(7)-Ag(20)-Ag(4)-Ag(14) | -0.0057 |
| Ag(7)-Ag(20)-Ag(4)-Ag(18) | -125.2723 |
| Ag(13)-Ag(12)-Ag(20)-Ag(4) | -125.2639 |
| Ag(13)-Ag(12)-Ag(20)-Ag(7) | 0.0007 |
| Ag(18)-Ag(12)-Ag(20)-Ag(4) | -0.0004 |
| Ag(18)-Ag(12)-Ag(20)-Ag(7) | 125.2642 |
| Ag(13)-Ag(8)-Ag(19)-Ag(10) | 109.4677 |
| Ag(13)-Ag(8)-Ag(19)-Ag(7) | -0.0062 |
| Ag(21)-Ag(8)-Ag(19)-Ag(10) | 0.0109 |
| Ag(21)-Ag(8)-Ag(19)-Ag(7) | -109.463 |
| Ag(13)-Ag(8)-Ag(21)-Ag(5) | 0.0018 |
| Ag(13)-Ag(8)-Ag(21)-Ag(11) | -125.2563 |
| Ag(19)-Ag(8)-Ag(21)-Ag(5) | 125.2513 |
| Ag(19)-Ag(8)-Ag(21)-Ag(11) | -0.0068 |
| Ag(5)-Ag(13)-Ag(8)-Ag(21) | -0.0018 |
| Ag(5)-Ag(13)-Ag(8)-Ag(19) | -125.2565 |
| Ag(12)-Ag(13)-Ag(8)-Ag(21) | 125.264 |
| Ag(12)-Ag(13)-Ag(8)-Ag(19) | 0.0093 |
| Ag(17)-Ag(11)-Ag(16)-Ag(10) | -125.2642 |
| Ag(17)-Ag(11)-Ag(16)-Ag(3) | -0.0066 |
| Ag(21)-Ag(11)-Ag(16)-Ag(10) | 0.0127 |
| Ag(21)-Ag(11)-Ag(16)-Ag(3) | 125.2704 |
| Ag(1)-Ag(18)-Ag(12)-Ag(13) | -0.009 |
| Ag(1)-Ag(18)-Ag(12)-Ag(20) | -125.2634 |
| Ag(4)-Ag(18)-Ag(12)-Ag(13) | 125.2548 |
| Ag(4)-Ag(18)-Ag(12)-Ag(20) | 0.0004 |
| Ag(18)-Ag(12)-Ag(13)-Ag(5) | 0.0098 |
| Ag(18)-Ag(12)-Ag(13)-Ag(8) | -109.4703 |
| Ag(20)-Ag(12)-Ag(13)-Ag(5) | 109.4736 |
| Ag(20)-Ag(12)-Ag(13)-Ag(8) | -0.0065 |
| Ag(17)-Ag(11)-Ag(21)-Ag(5) | 0.0039 |
| Ag(17)-Ag(11)-Ag(21)-Ag(8) | 109.475 |
| Ag(16)-Ag(11)-Ag(21)-Ag(5) | -109.4762 |
| Ag(16)-Ag(11)-Ag(21)-Ag(8) | -0.005 |
| Ag(3)-Ag(17)-Ag(11)-Ag(21) | -125.2605 |
| Ag(3)-Ag(17)-Ag(11)-Ag(16) | 0.0066 |
| Ag(9)-Ag(17)-Ag(11)-Ag(21) | 0.0008 |
| Ag(9)-Ag(17)-Ag(11)-Ag(16) | 125.2679 |
| Ag(16)-Ag(10)-Ag(24)-Ag(2) | -0.0127 |
| Ag(16)-Ag(10)-Ag(24)-Ag(7) | -125.2633 |
| Ag(19)-Ag(10)-Ag(24)-Ag(2) | 125.2535 |
| Ag(19)-Ag(10)-Ag(24)-Ag(7) | 0.0029 |
| Ag(8)-Ag(19)-Ag(10)-Ag(16) | -0.0032 |
| Ag(8)-Ag(19)-Ag(10)-Ag(24) | -125.2751 |
| Ag(7)-Ag(19)-Ag(10)-Ag(16) | 125.269 |
| Ag(7)-Ag(19)-Ag(10)-Ag(24) | -0.0029 |
| Ag(11)-Ag(16)-Ag(10)-Ag(19) | -0.0086 |
| Ag(11)-Ag(16)-Ag(10)-Ag(24) | 109.4705 |
| Ag(3)-Ag(16)-Ag(10)-Ag(19) | -109.4749 |
| Ag(3)-Ag(16)-Ag(10)-Ag(24) | 0.0042 |
| Ag(1)-Ag(23)-Ag(9)-Ag(15) | 0.0017 |
| Ag(1)-Ag(23)-Ag(9)-Ag(17) | -125.2571 |
| Ag(5)-Ag(23)-Ag(9)-Ag(15) | 125.2601 |
| Ag(5)-Ag(23)-Ag(9)-Ag(17) | 0.0012 |
| Ag(3)-Ag(17)-Ag(9)-Ag(15) | -0.0082 |
| Ag(3)-Ag(17)-Ag(9)-Ag(23) | 109.4588 |
| Ag(11)-Ag(17)-Ag(9)-Ag(15) | -109.4704 |
| Ag(11)-Ag(17)-Ag(9)-Ag(23) | -0.0034 |
| Ag(1)-Ag(15)-Ag(9)-Ag(17) | 125.2665 |
| Ag(1)-Ag(15)-Ag(9)-Ag(23) | -0.0017 |
| Ag(6)-Ag(15)-Ag(9)-Ag(17) | 0.0009 |
| Ag(6)-Ag(15)-Ag(9)-Ag(23) | -125.2674 |
| Ag(14)-Ag(6)-Ag(22)-Ag(3) | -125.263 |
| Ag(14)-Ag(6)-Ag(22)-Ag(2) | -0.008 |
| Ag(15)-Ag(6)-Ag(22)-Ag(3) | 0.0075 |
| Ag(15)-Ag(6)-Ag(22)-Ag(2) | 125.2625 |
| Ag(14)-Ag(6)-Ag(15)-Ag(1) | -0.0022 |
| Ag(14)-Ag(6)-Ag(15)-Ag(9) | 109.4764 |
| Ag(22)-Ag(6)-Ag(15)-Ag(1) | -109.4791 |
| Ag(22)-Ag(6)-Ag(15)-Ag(9) | -0.0005 |
| Ag(15)-Ag(6)-Ag(14)-Ag(4) | -0.0058 |
| Ag(15)-Ag(6)-Ag(14)-Ag(2) | -125.2585 |
| Ag(22)-Ag(6)-Ag(14)-Ag(4) | 125.2607 |
| Ag(22)-Ag(6)-Ag(14)-Ag(2) | 0.008 |
| Ag(13)-Ag(5)-Ag(23)-Ag(1) | -0.0035 |
| Ag(13)-Ag(5)-Ag(23)-Ag(9) | -109.4721 |
| Ag(21)-Ag(5)-Ag(23)-Ag(1) | 109.4721 |
| Ag(21)-Ag(5)-Ag(23)-Ag(9) | 0.0034 |
| Ag(11)-Ag(21)-Ag(5)-Ag(13) | 125.2646 |
| Ag(11)-Ag(21)-Ag(5)-Ag(23) | -0.006 |
| Ag(8)-Ag(21)-Ag(5)-Ag(13) | -0.0018 |
| Ag(8)-Ag(21)-Ag(5)-Ag(23) | -125.2724 |
| Ag(12)-Ag(13)-Ag(5)-Ag(21) | -125.2745 |
| Ag(12)-Ag(13)-Ag(5)-Ag(23) | -0.0036 |
| Ag(8)-Ag(13)-Ag(5)-Ag(21) | 0.0018 |
| Ag(8)-Ag(13)-Ag(5)-Ag(23) | 125.2728 |
| Ag(1)-Ag(18)-Ag(4)-Ag(14) | -0.0087 |
| Ag(1)-Ag(18)-Ag(4)-Ag(20) | 125.2605 |
| Ag(12)-Ag(18)-Ag(4)-Ag(14) | -125.2696 |
| Ag(12)-Ag(18)-Ag(4)-Ag(20) | -0.0004 |
| Ag(18)-Ag(4)-Ag(14)-Ag(6) | 0.0113 |
| Ag(18)-Ag(4)-Ag(14)-Ag(2) | 109.4706 |
| Ag(20)-Ag(4)-Ag(14)-Ag(6) | -109.4614 |
| Ag(20)-Ag(4)-Ag(14)-Ag(2) | -0.0021 |
| Ag(17)-Ag(3)-Ag(22)-Ag(6) | -0.0148 |
| Ag(17)-Ag(3)-Ag(22)-Ag(2) | -109.4771 |
| Ag(16)-Ag(3)-Ag(22)-Ag(6) | 109.4648 |
| Ag(16)-Ag(3)-Ag(22)-Ag(2) | 0.0025 |
| Ag(22)-Ag(3)-Ag(17)-Ag(9) | 0.0152 |
| Ag(22)-Ag(3)-Ag(17)-Ag(11) | 125.2672 |
| Ag(16)-Ag(3)-Ag(17)-Ag(9) | -125.2585 |
| Ag(16)-Ag(3)-Ag(17)-Ag(11) | -0.0066 |
| Ag(10)-Ag(24)-Ag(2)-Ag(14) | -109.4689 |
| Ag(10)-Ag(24)-Ag(2)-Ag(22) | 0.0162 |
| Ag(7)-Ag(24)-Ag(2)-Ag(14) | -0.0138 |
| Ag(7)-Ag(24)-Ag(2)-Ag(22) | 109.4713 |
| Ag(5)-Ag(23)-Ag(1)-Ag(15) | -125.2605 |
| Ag(5)-Ag(23)-Ag(1)-Ag(18) | 0.0043 |
| Ag(9)-Ag(23)-Ag(1)-Ag(15) | -0.0017 |
| Ag(9)-Ag(23)-Ag(1)-Ag(18) | 125.2631 |
| Ag(15)-Ag(1)-Ag(18)-Ag(4) | 0.0007 |
| Ag(15)-Ag(1)-Ag(18)-Ag(12) | 109.4719 |
| Ag(23)-Ag(1)-Ag(18)-Ag(4) | -109.4693 |
| Ag(23)-Ag(1)-Ag(18)-Ag(12) | 0.0019 |
| Ag(6)-Ag(15)-Ag(1)-Ag(18) | 0.0047 |
| Ag(6)-Ag(15)-Ag(1)-Ag(23) | 125.2728 |
| Ag(9)-Ag(15)-Ag(1)-Ag(18) | -125.2664 |
| Ag(9)-Ag(15)-Ag(1)-Ag(23) | 0.0017 |
